# The transcriptional landscape of the murine middle ear epithelium *in vitro*

**DOI:** 10.1101/800987

**Authors:** Apoorva Mulay, Md Miraj K Chowdhury, Cameron James, Lynne Bingle, Colin D Bingle

**Author notes:** Cedars-Sinai Medical Centre 8700 Beverly Blvd, AHSP, Los Angeles CA 90048. Corresponding author: Colin Bingle PhD, Department of Infection, Immunity and Cardiovascular Disease, The Medical School, University of Sheffield, Sheffield, S10 2RX, UK.

## Abstract

Otitis media (OM) is the most common paediatric disease and leads to significant morbidity. Although understanding of underlying disease mechanisms is hampered by complex pathophysiology, it is clear that epithelial abnormalities underpin the disease. The mechanisms underpinning epithelial remodelling in OM remain unclear. We recently described a novel in vitro model of mouse middle ear epithelial cells (mMEECs) that undergoes mucociliary differentiation into the varied epithelial cell populations seen in the middle ear cavity. We now describe genome wide gene expression profiles of mMEECs as they undergo differentiation. We compared the gene expression profiles of original (uncultured) middle ear cells, confluent cultures of undifferentiated cells (day 0 of ALI) and cells that had been differentiated for 7 days at an ALI. >5000 genes were differentially expressed among the three groups of cells. Approximately 4000 genes were differentially expressed between the original cells and day 0 of ALI culture. The original cell population was shown to contain a mix of cell types, including contaminating inflammatory cells that were lost on culture. Approximately 500 genes were upregulated during ALI induced differentiation. These included some secretory genes and some enzymes but most were associated with the process of ciliogenesis. Our in vitro model of differentiated murine middle ear epithelium exhibits a transcriptional profile consistent with the mucociliary epithelium seen within the middle ear. Knowledge of the transcriptional landscape of this epithelium will provide a basis for understanding the phenotypic changes seen in murine models of OM.

## Introduction

Otitis media (OM), a group of inflammatory diseases of the middle ear, is the leading cause of paediatric surgery and the most frequent reason for the prescription of antibiotics [1,2]. It can have both acute and chronic (recurrent) presentations the consequences of which may be life-long. Over 80% of children develop at least one incidence of OM by three years of age and over 700 million cases occur world-wide per year [3].

Children with recurrent episodes of OM are at risk of developing hearing loss and it is estimated that, globally, over 100 million people have hearing loss due to OM. Consequently, the disease is a major paediatric clinical problem that produces significant morbidity and quality-of-life issues across the world, with the major burden being seen in children in developing countries [3].

The middle ear epithelium and its secretions, are involved in maintaining homeostasis and sterility within the middle ear cavity (MEC). Multiple host defence proteins have been shown to help protect the middle ear cavity. For example, we recently identifed BPIFA1 as an abundant secretory protein produced by the middle ear epithelium and showed that loss of the gene exacerbated disease severity in an established model of OM [4]. Furthermore, loss of the gel forming mucin, MUC5B has also been shown to cause OM [5]. OM can be considered to be a disease of the middle ear epithelium. The epithelial lining of the middle ear cavity varies according to the location [6,7]. The attic, or epitympanum, is lined by squamous epithelium while the middle ear proper is lined by cuboidal epithelium and the hypotympanum by ciliated columnar epithelium. Phenotypic changes throughout the middle ear epithelium are key to the pathophysiology of OM [7,8]. Current knowledge suggests that OM is caused by an unrestrained response by the middle ear epithelium to an exogenous trigger, often a pathogen [9,10]. This is associated with excess production of secretory proteins from the abnormal epithelium, which along with inflammatory cells, produce middle ear exudate [1]. The mechanisms underpinning the epithelial remodelling remain unclear but result in the development of an abnormal mucociliary epithelium that contributes to the development of characteristic exudates.

The ability to identify the function of different cell types and their products within the middle ear has limited our understanding of phenotypic changes underpinning OM development. Research into the pathogenesis of OM is limited because of difficulties in accessing appropriate samples, at an early stage in the disease process. Most studies have involved sampling the middle ear fluid [11] or been restricted to whole animal challenge studies [12-15]. We recently described the establishment of an air liquid interface (ALI) culture system to model the mouse middle ear epithelium *in vitro* [16,17]. We showed that a “basal cell like” population of middle ear epithelial cells underwent differentiation during ALI culture to generate a complex mucosal tissue that had characteristics of the native middle ear epithelium [16] We now describe genome wide gene expression profiles of these cells as they undergo differentiation to provide an understanding of the transcriptional landscape of this complex epithelium.

## Materials and Methods

### Ethics statement

Humane care and animal procedures were carried out in accordance to the appropriate UK Home Office Project licence. 8-10 week old C57BL/6 mice housed in individually ventilated cages (Techniplast UK Ltd) under specific pathogen free (SPF) conditions were obtained from MRC Harwell, UK.

### Isolation and differentiation of middle ear epithelial cells at air liquid interface

The protocol for primary culture and differentiation of mouse middle ear epithelial cells (mMEECs) has been described in detail previously [16,17]. Five or six mice (10-12 bullae) were used for each cell isolation. Following a differential adherence step which removes contaminating fibroblasts, the original cell population contains a mix of cell types including differentiated epithelial cells, multipotent “basal cell-like” epithelial cells and inflammatory cells. 1 × 10^4^ of these “original” cells were plated on to rat-tail collagen I coated 24-well, 0.4μm pore sized transparent PET (Polyethylene Terephthalate) membranes in the presence of 10 μM of Rho Kinase inhibitor, Y-27632 dihydrochloride (ROCKi, Tocris bioscience, Cat-1254). Cells were cultured to confluence in submerged culture and subsequently grown at the ALI for up to 7 days. Cells were lysed in 250 µL of Trizol reagent (Sigma-Aldrich, Cat No-T2494) for RNA extraction at ALI Day 0, Day 3, and Day 7. Figure 1A gives a brief overview of the complete cell culture system.

**Figure 1:**
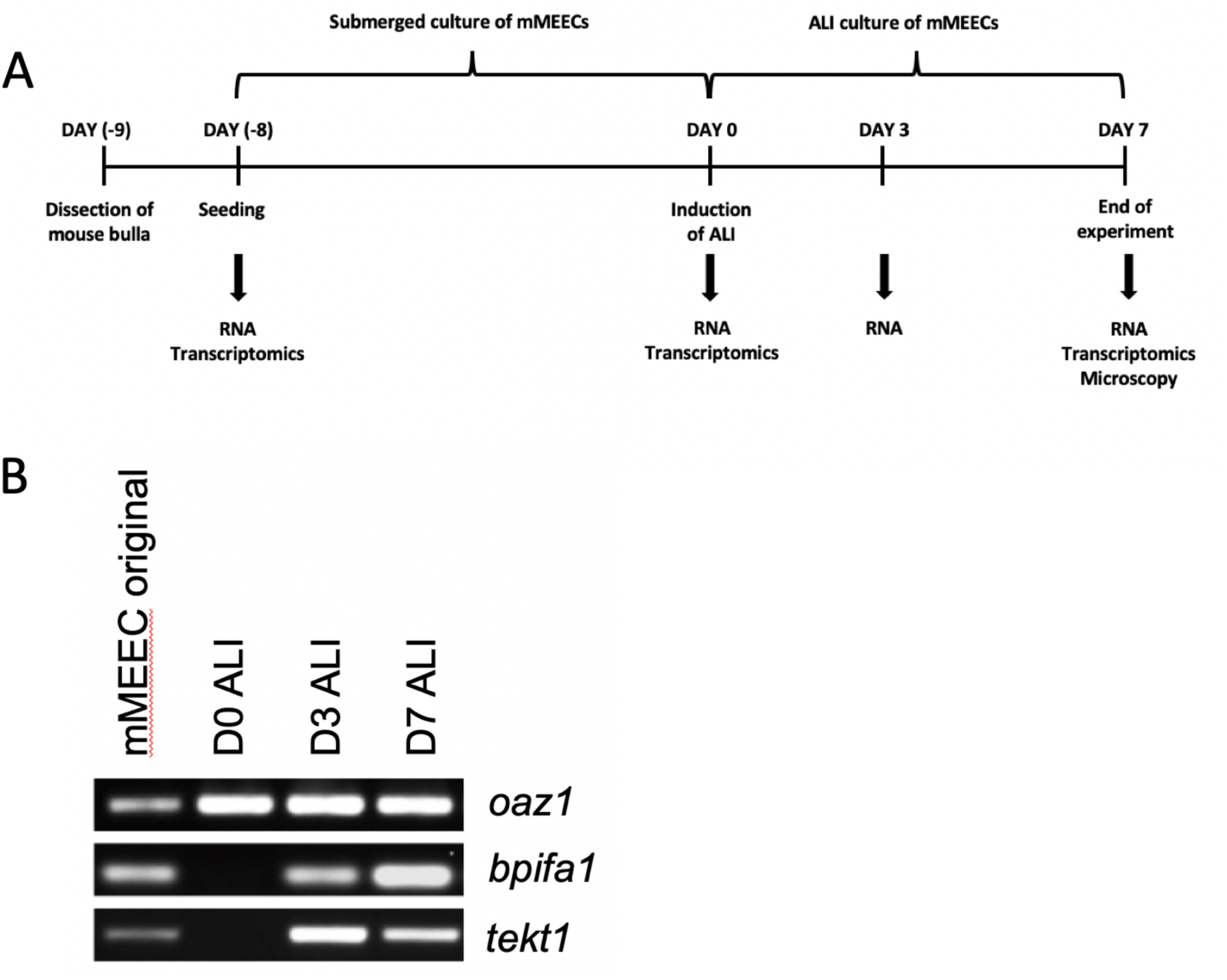
Culture of mouse middle ear epithelial cells for transcriptional analysis. **A.** Schematic timeline for culture experiments mMECs Middle ear epithelial cells isolated from dissected bullae (day -9) and cultured in transewells in submerged culture till confluence (day -8 to day 0), before ALI was induced. Samples for transcriptional analysis were collected at seeding (original cells), day 0, day 3 and day 7. **B**. End-point RT-PCR showing expression of Oaz1, Bpifa1 and Tekt1 in mMEC original cells isolated from the middle ear cavity, ALI day 0, 3 and 7 cells

### RNA extraction and Reverse transcription PCR (RT-PCR)

For end-point RT-PCR, total RNA was extracted from freshly isolated mMEECs before seeding (original) and cells at ALI days 0, 3 and 7 lysed in Trizol. RNA yield was determined using NanoDrop-1000 (Thermoscientific). Residual genomic DNA was digested by DNase I treatment (Promega, Cat No-M6101) and 200ng of RNA was reverse transcribed using AMV Reverse Transcriptase (Promega, Cat No-M9004). RT-PCR was performed with 1µl of template cDNA and Maxima Hot Start Green PCR Master Mix (ThermoFisher Scientific, Cat No-K1061). The cycling conditions were: 95°C for 5 minutes; denaturation: 94°C for 1 minute (25-35 cycles); annealing: 60°C for 1 minute; extension: 72°C for 1 minute; final extension: 72°C for 7 minutes (MJ Research PTC-200). The primer pairs used are as follows. Bpifa1F, ACAGAGGAGCCGACGTCTAA; Bpifa1R, CCAAGAAAGCTGAAGGTTC;Tekt1 F, CAGTGCGAAGTGGTAGACG; Tekt1R, TTCACCTGGATTTCCTCCTG; Oaz1F, ACAGAGGAGCCGACGTCTAA; Oaz1R, CCAAGAAAGCTGAAGGTTC; Spata18F, GCAATGCAGTCCTTAGAGCC; Spata18R,CATTACTGGTCGCACGGAC; Dynlrb2F, CCACAGGCGCGATGACAG;Dynlrb2R,ACTCACATGGGTTCTGAATGACA;Cdhr3F, AGGTGGAAAGGCCCATTAAC; Cdhr3R, AGTCGTAGAAGGGCATCAGG. The amplified PCR products were run on a 2% agarose gel containing 0.5µg/ml ethidium bromide (Dutscher scientific, Cat No-4905006) and bands visualised using a Biorad ChemiDoc™ XRS+.

### Clariom S mouse microarray

Samples of total RNA were checked for quantity and quality using the Nanodrop and Agilent Bionalyser 600. Subsequently 200ng was prepared for analysis on the Clariom S mouse GeneChip (Thermo Fisher) according to the manufacturers’ instructions. Briefly mRNA was converted to double stranded cDNA with the incorporation of a T7 polymerase binding site at the 3’ end of the RNA molecule. Antisense RNA was generated by utilization of the T7 polymerase. This was purified using a magnetic bead process and further quantified on the Nanodrop. 15ug of the aRNA was taken forward to generate a sense DNA strand. This was fragmented and end labelled with biotin before being incorporated into a hybridization solution and incubated with the GeneChip. Following post hybridisation washing using the fluidics station a fluorescent signal corresponding to hybridization of the labelled material to the oligonucleotide probes on the chip was achieved using a streptavidin-phycoerythrein cocktail. Gene chips were scanned on the Gene Chip 7000G scanner and the images collected as CEL files.

### Bioinformatic analysis

Microarray data in CEL files was analysed using Affymetrix Expression Console. Gene level summary data were obtained using the RMA summarization method. Differentially expressed genes were identified using limma [18] with the criteria of absolute fold change > 2 and limma adjusted p value < 0.05. Batch effect was included in the statistical models. p values were adjusted with Benjamini and Hochberg method [19]. A gene can have more than one probe sets. If a gene has both up- and down-regulated probe sets, it is removed from the list of differentially expressed genes. Multidimensional scaling plot were generated using limma. Gene ontology analyses were performed using clusterProfiler [20].

## Results

We previously showed that mMEECs undergo a process of mucociliary differentiation over a period of 14 days when cultured at the ALI. In this study we used cells that were isolated from mouse bullae and cultured for 7 days at the ALI (shown schematically in Figure 1A). We used end point PCR to confirm that the cells switched on genes associated with differentiated secretory (*bpifa1*) and ciliated cells (*tekt1*) within three days of establishment of ALI culture (figure 1B). We then used gene expression array analysis to compare the gene expression profiles of matched, triplicate samples of original (uncultured) middle ear cells, confluent cultures of undifferentiated cells (day 0 of ALI) and cells that had been differentiated for 7 days at the ALI (Table 1). The original cells used to initiate the cultures would be expected to contain a mixture of cells including different epithelial cells along with inflammatory and blood cells recovered from the tissue during isolation.

**Table 1.**
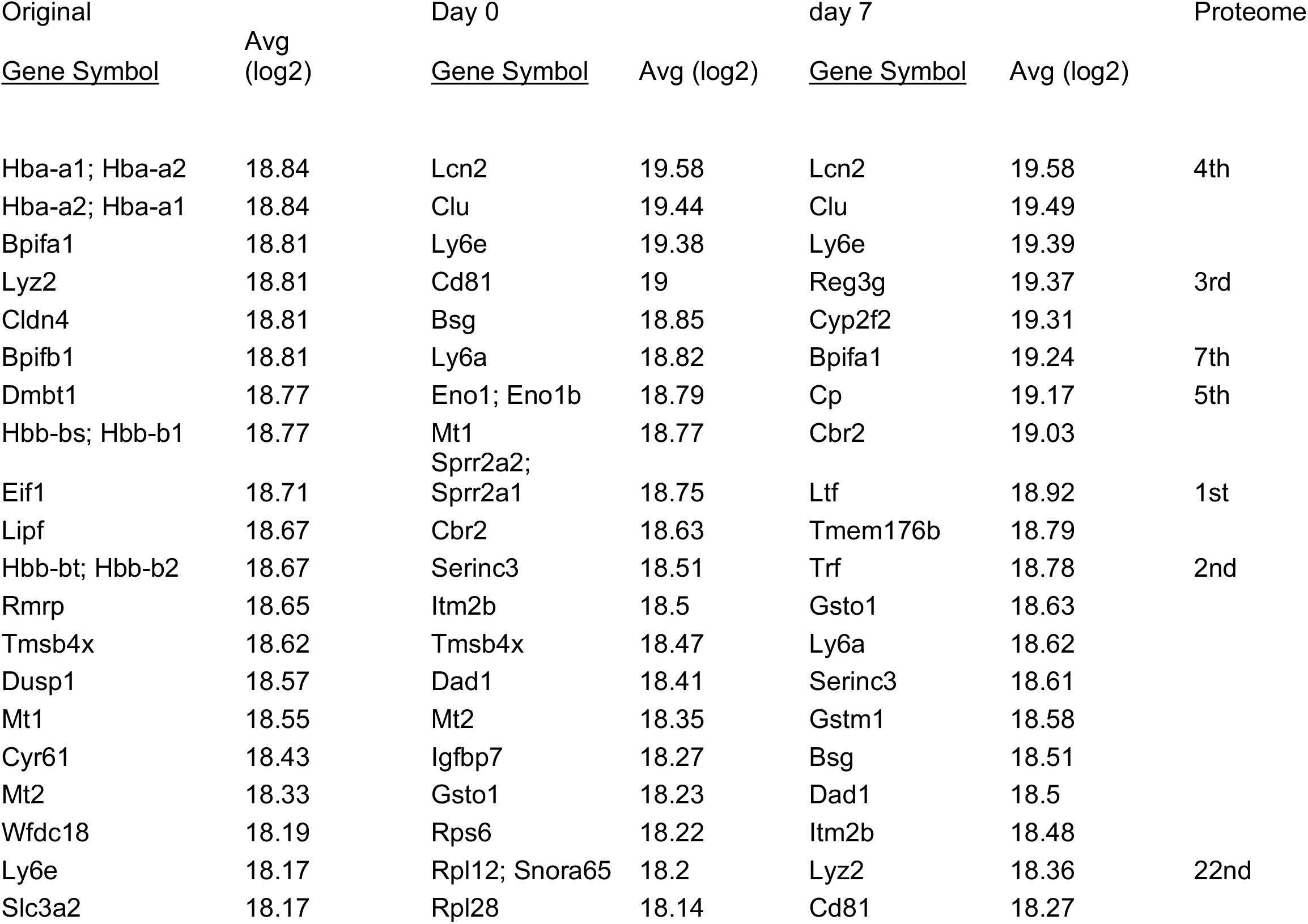
The top 20 genes from each set of samples are listed in descending level of expression. Data was log transformed. The proteome column shows the ranking of each secreted protein in the day 14 secreted proteome of ALI MEECs cells taken from Mulay *et al*, 2016 [16].

Principal Component Analysis (Figure 2A) showed that the three different sample groups were clustered together, although the original cell sample exhibited the greatest variability. The 20 most highly expressed at each time point are listed in Table 1. In the original cells four of these genes represented haemaglobin genes and the three top secretory protein gene were *Bpifa1, Lyz2* and *Bpifb1*. At Day 0 of culture all but one of the top 20 genes were structural, with the exception of *Lcn2*. By day 7 of ALI culture the most highly expressed genes included multiple secretory protein genes including, *Lcn2, Reg3g, Bpifa1, Cp, Ltf* and *Tf*. This expression data is in keeping with our previous proteomic data which identifed these as being amongst the most abundant proteins in apical secretions from ALI cells (Table 1) [16].

**Figure 2:**
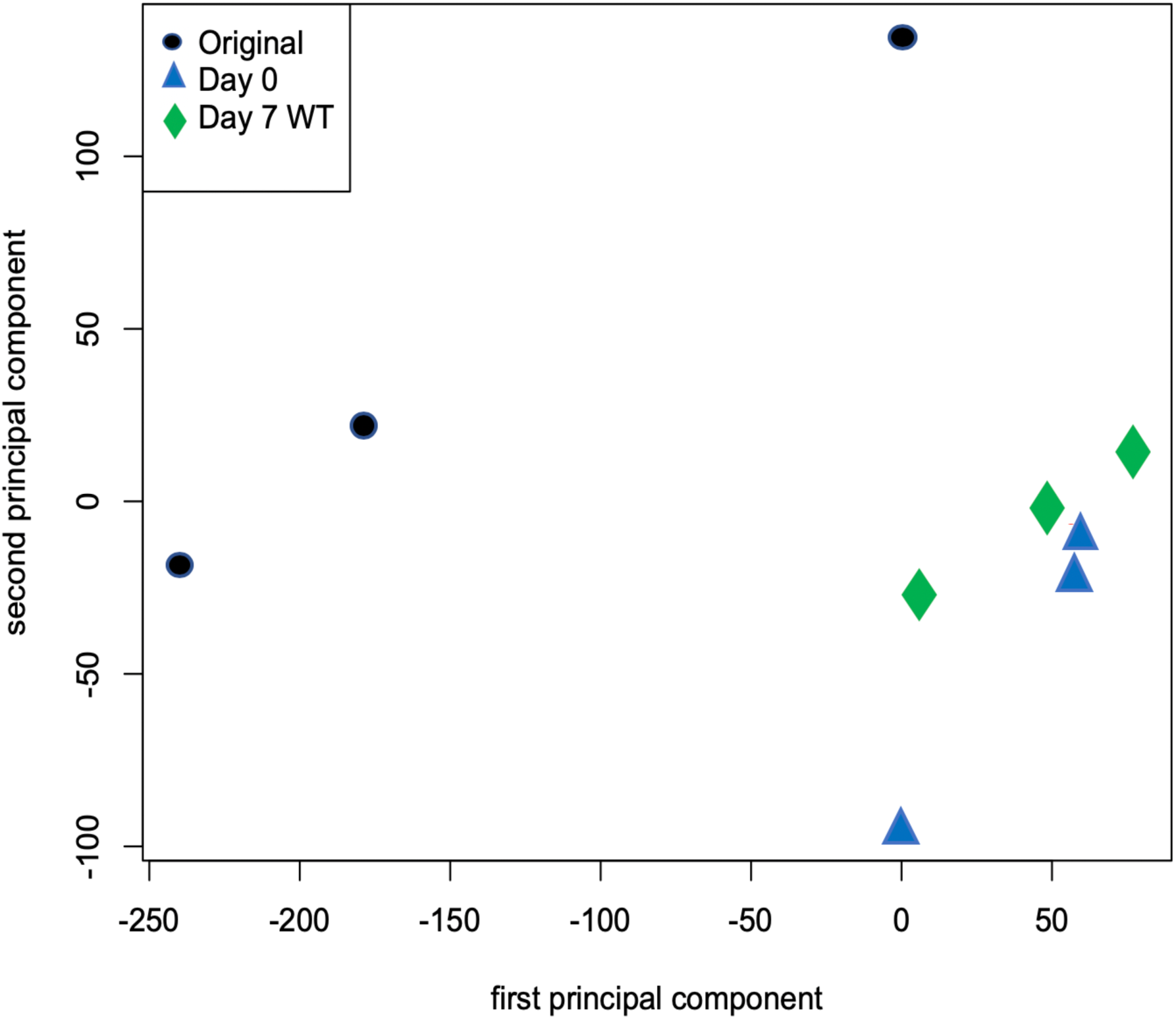
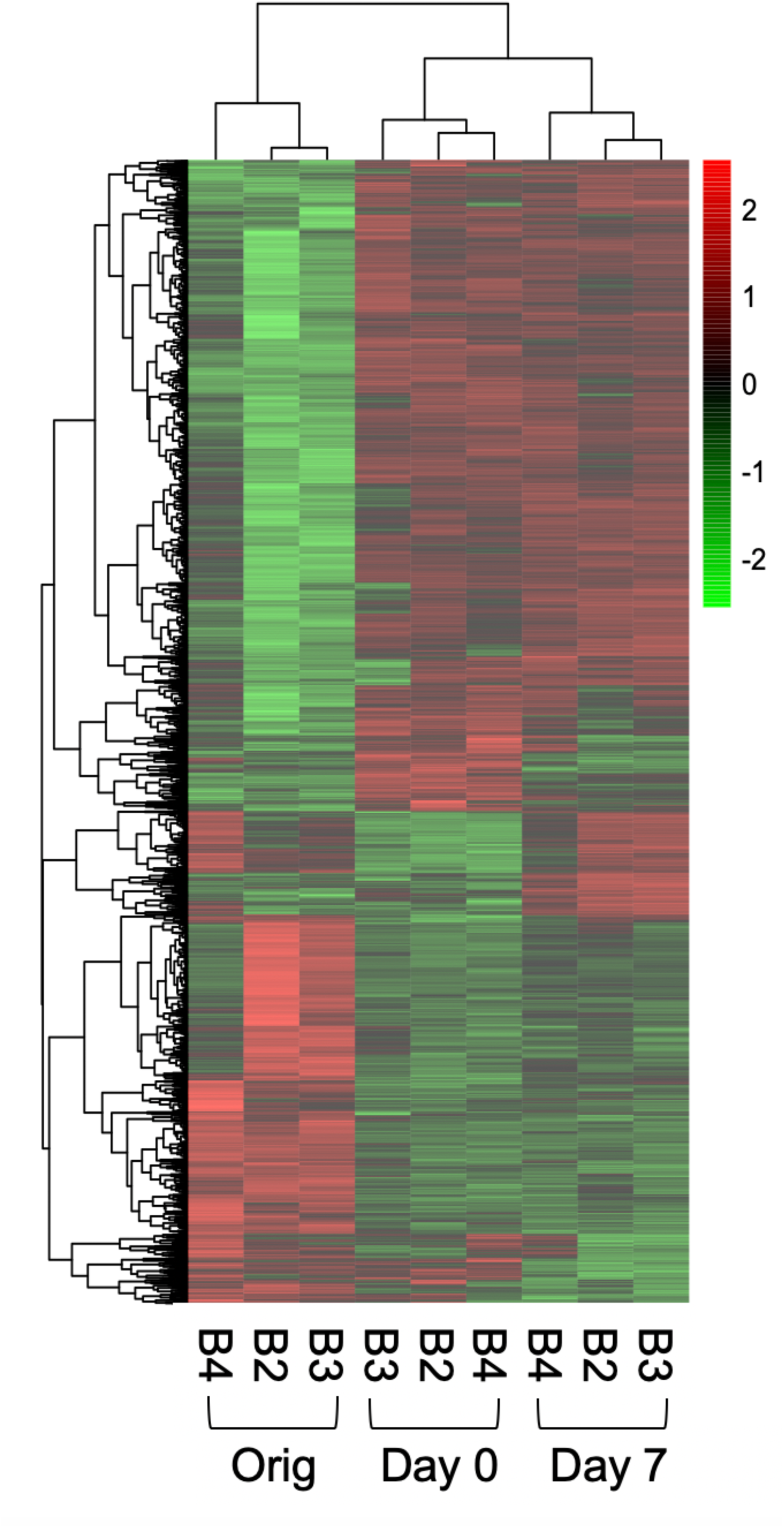
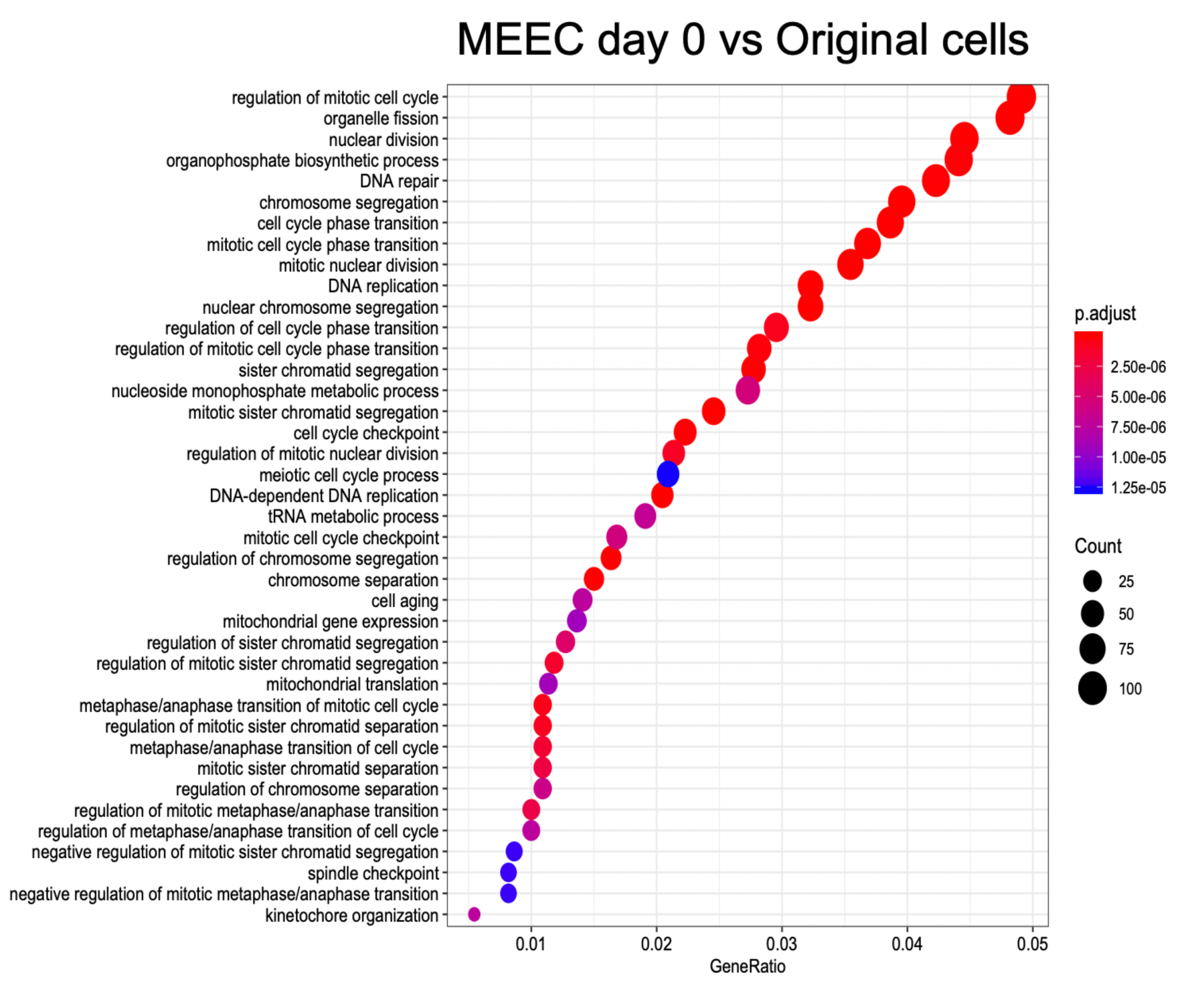
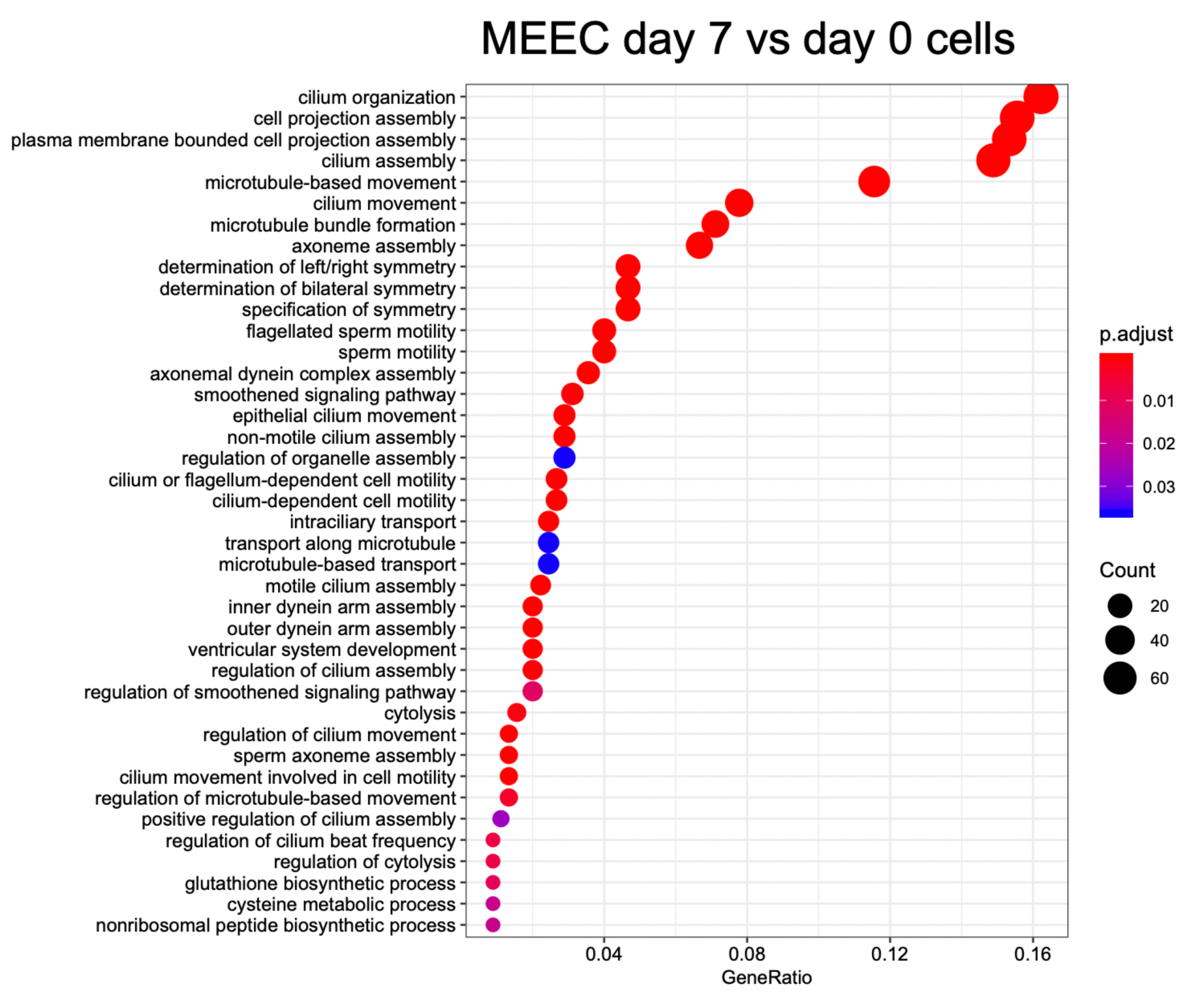
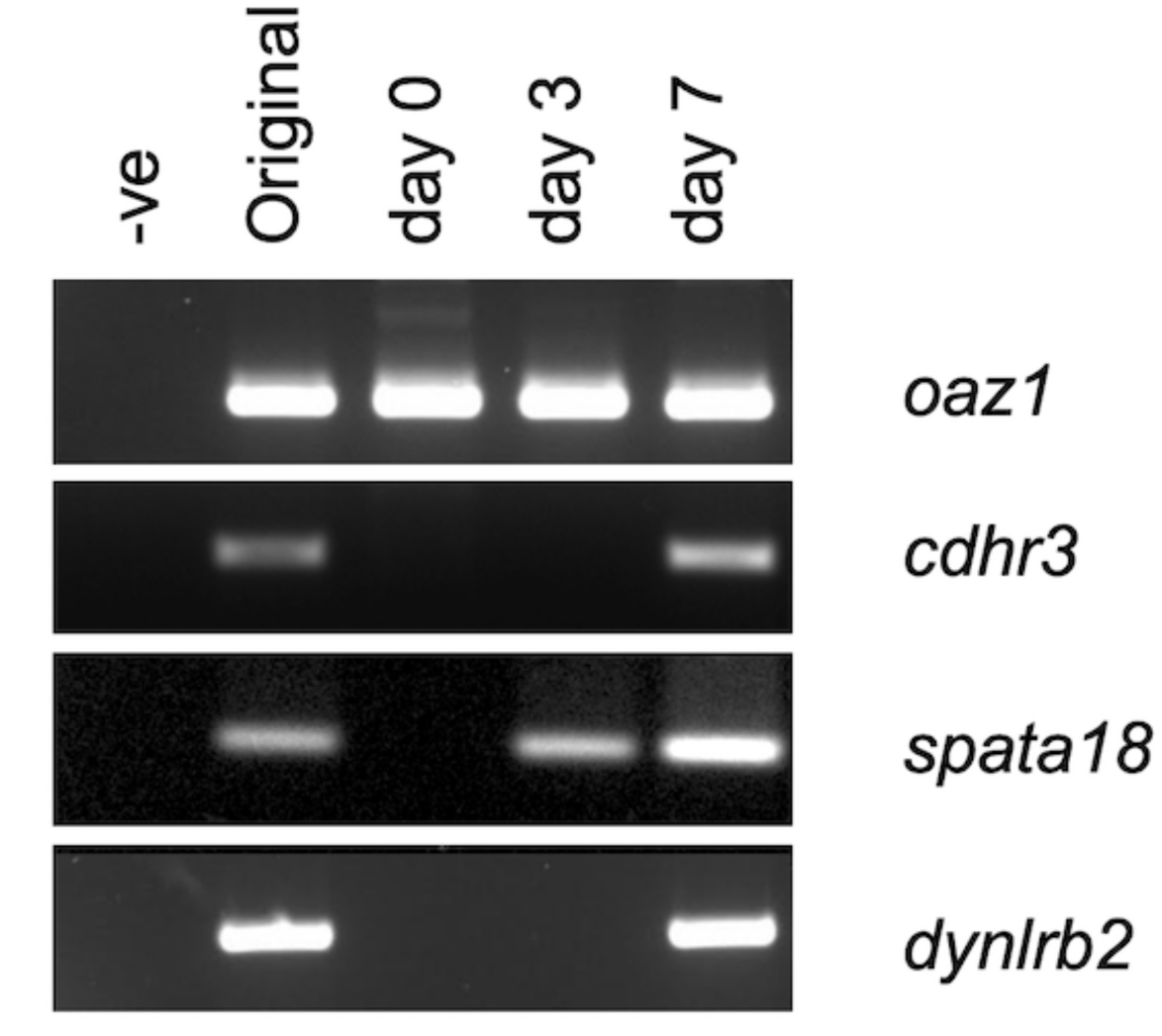
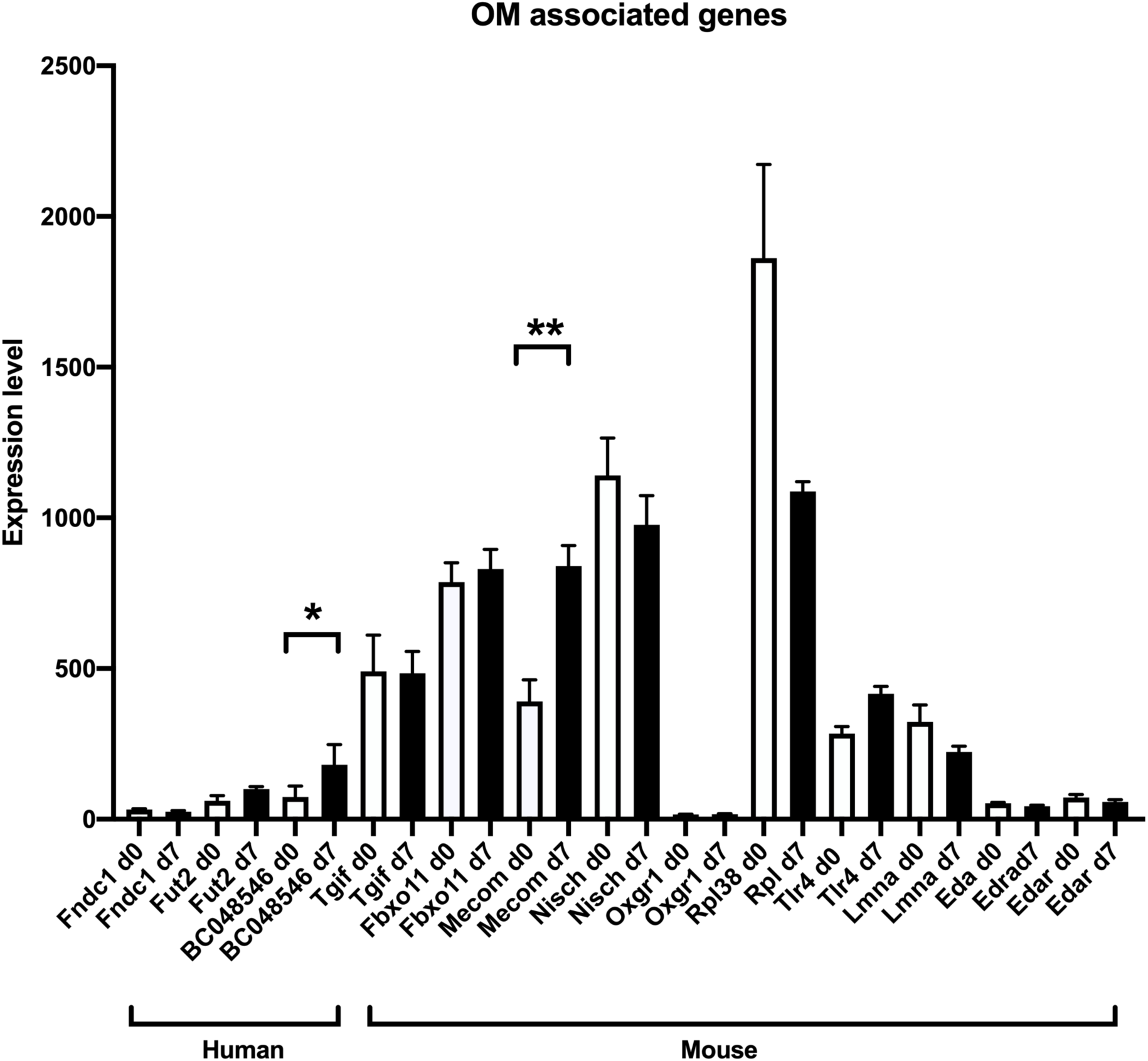
Genome wide transcriptional analysis of differentiating mMEECs. **A.** PCA analysis was used to show the relatedness of the different samples used in the study. **B.** The heatmap shows differentially expressed genes identified by Limma. Genes used in the analysis were >2 fold differentially expressed. **C**,**D.** Gene ontogeny analysis of the most highly differentially expressed genes are displayd for day 0 vs original cells (**C)** and Day 7 vs day 0 (**D**). **E.** End-point RT-PCR showing expression of *Oaz1, Cdhr3, Spata18* and *Dynlrb2* in cDNA from mMEC original cells isolated from the middle ear cavity, ALI day 0, 3 and 7 cells. Negative control samples contained no cDNA. **F**. Microarray derived data of relative expression of candidate OM associated genes in day 0 and day 7 cells. * P >0.05; ** P >0.01 using log transformed data. Human and mouse candidate genes are identified.

Following Robust MuitiArray Average (RMA) normalisation differential gene expression was undertaken using limma [21] This analysis (Figure 2B) revealed that a total of 5261 genes were differentially expressed amongst the three different groups with an absolute fold change of >2 and a limma adjusted p value of <0.05. The heat map clearly shows that samples from the different time points group together but also shows some level of variability between individual samples. When comparing different time points 1580 genes were significantly more highly expressed in the original cells compared to the Day 0 cultures whereas 2400 genes were more highly expressed in the Day 0 cells compared to the original cells (Table S1). The most highly differentially expressed genes in the original samples were, *Lipf, Hbb-bs, Hbb-bt, Bpifb1* and *Lyz1* whereas in the Day 0 cells they were *Arg1, Sorbs2, Srd5a1, Vsnl1* and *Lox.* When Day 0 cells were compared to Day 7 ALI cells many fewer genes were differentially expressed. 489 genes were up regulated as the cells differentiated whereas expression of 385 genes were significantly reduced (Table S2). The most highly differentially expressed genes in the Day 7 samples were, *Lyz2, Aldh1a1, Spata18, Cdhr3* and *Lrrc34*, whereas in the Day 0 cells they were *Ppbp (Cxcl7), Lgals1 (Galectin1), Il1a, Ndufa412* and *2610528A11Rik (Gpr15l).* Comparisons between the original cells and the Day 7 ALI cells showed that 2408 genes were up regulated whereas 1215 genes were down regulated (Table S3). The most highly differentially expressed genes in the Day 7 samples were, *Fetub, Sorbs2, Rgs5, Serpina7* and *Kcnj16*, whereas in the original cells they were *Lipf, Bpifb1, Hbb-bs, Hbb-bt*, and *Hba*-a2.

To gain more useful information from the differential gene expression data sets we subjected them to gene ontology analysis using clusterProfiler [20]. For this analysis we were most interested in understanding the processes that occurred as epithelial cells became confluent during the submerged culture period and what happened when the cells underwent differentiation. With this in mind we focused our analysis on identifying genes upregulated as the cells became confluent and also on those that were upregulated as the cells differentiated. The top biological processes enriched as the cells reached confluence at Day 0 included those associated with cell division, cell migration and cell activation (Figure 2C). In the Day 7 cells the majority of pathways identified were associated with aspects of ciliogenesis or ciliary function (Figure 2D). Close inspection of the differential gene list from this time point allowed the identification of many genes that are shown to be associated with cilia, including *Spata18, Cdhr3, Lrrc34, Dynlrb2, Lrguk, Dnah6, Ccdc67, Dhrs9, Spef2*, and *ccdc146. Tekt1* the gene we use as a ciliated cell specific marker in our end-point PCR reactions was the 26^th^ most differentially expressed in this list (Table S2). We used end-point PCR with RNA extracted from a different batch of cells to confirm that *Spata18, Cdhr3* and *dynlrb2* were upregulated during the process of differentiation (Figure 2E).

### Expression of OM associated genes in the in vitro mouse middle ear epithelium

One of the potential uses of our *in vitro* model is that is may be useful for the study of genes associated with OM. Multiple genetic association studies have been undertaken to identify human genes associated with OM [22-24]. Additionally, loss of function of multiple genes have been shown to lead to the development of OM-like phenotypes in mice [25,26]. Expression of three well validate human OM associated genes *Fndc1 [27], Fut2* [28] and *A2ml1* (*BC048546*) [29] was detected in the mMMECs with the highest expression being for *BC048546* in the day 7 differentiated cells (Figure 2F). Expression of four OM associated genes identified through ENU mutagenesis screening: *Tgif1 [30], Fxbxo11* [31], *Mecom* [32] and *Nisch* [33} is clearly seen in both the undifferentiated and differentiated cells, with *Mecom* (*Evi1*) the gene mutated in the *Junbo*^+*/-*^ OM mouse model [4,32] being most differentially expressed during differentiation. The expression of a further, representative group of genes associated with murine OM was shown to be variable with *Oxgr1* [34] being essentially not expressed in the cells whereas *Rpl38* [35] was the most highly expressed. This simple analysis shows that, although there was not clear differentiation associated changes in many of these genes, this data set will be useful to understand something about the epithelial expression of OM candidate genes.

## Discussion

The mucosal lining of the middle ear cavity covers the entire surface and varies according to the location [6,7]. Cellular morphology can be simple squamous, cuboidal or columnar depending on the location and can have tracts of ciliated cells interspersed with secretory cells. We recently described a method to differentiate primary mMEECs at an air liquid interface. We could show that the differentiated cultures took on a morphology similar to that seen *in situ* with tracts of elevated columnar mucociliary epithelium that contained both ciliated (FOXJ1^+ve^) and goblet (MUC5B^+ve^) cells alongside cuboidal (BPIFA1^+ve^) cells [16]. This model system overcomes a major limitation of previous middle ear epithelial cultures, namely the lack of differentiation into distinct epithelial cell types that are hallmarks of the mucociliary epithelium seen in the murine middle ear.

In this study we have used an unbiased expression array approach to better define the transcriptional signature of this *in vitro* murine middle ear epithelium. We chose to study a single time point of seven days of differentiation so as to capture the development of early stages of the mucociliary differentiation that occurs in this model. The transcriptional differences in the cultures at the three different times points were striking. The original cells, a cell population that had been depleted of fibroblasts (perhaps not completely, as judged by expression of *Col1a1* and *Col1a2* in the samples) by a differential adherence step, were made up of a mixture of cells from the middle ear. In this cell population *Bpifa1* was the most highly expressed secretory gene, followed by *Lcn2.* Unsurprisingly, this population of cells exhibited a strong differential gene expression signature for myeloid cells, including macrophages and neutrophils, represented by *S100a8, Ngp, Lilrb4a, Retnlg* and *Spi1*. Amongst the other highly expressed genes in the original cells were multiple Hemaglobin genes, potentially indicating that some blood contaminated the isolated cells. However, Hb genes have been shown to be expressed in multiple cells as well as in erythrocyte precursors [36-38]. There was also very high differential expression of *Lyz1* and *Lyz2* both of which are known to be expressed in both macrophages and structural lung cells [39]. The list also included some unexpected genes. For example, the top differentially expressed gene was gastric lipase (*Lipf*) a gene that has not previously shown to be expressed in the ear. Consistent with our previous data [16], we could show that *Bpifb1* was highly expressed in the RNA (the most highly expressed secretory gene) from the original cell population but was lost in the day 0 cells. Expression of this gene did not recover to the levels of the original after 7 days of differentiation. BPIFB1 is a marker of a population of goblet cells in the upper respiratory tract and nasopharynx of mice [40] and is present in human COME exudates [11]. Consistent with the gene expression data we did not detect BPIFB1 in apical secretions from MMECs [16]. We assume that the *Bpifb1* signal comes from a goblet cell population from the eustachian tube.

During the 7 day period when the cells were grown at the ALI the most striking signature seen across differentially expressed genes was one of cilia and ciliogenesis. Amongst the top differentially expressed genes were also some secretory protein genes. This observation confirms that the cultured cells underwent mucociliary differentiation. This gene signature is consistent with our previous IF studies which showed the presence of ciliated, secretory and basal cells in the cultures [16]. This *in vitro* data is also consistent with *in vivo* studies which have shown the presence of cuboidal epithelial cells expressing secretory proteins alongside regions of ciliated cells [6,7]. Although some of the ciliated cell signature genes are reasonably well studied such as; Spata18 [41] and Cdhr3 [42], none have been identified in the ear previously. The process multiciliogenesis is very complex and involves the concerted action of multiple pathways that regulate hundreds of genes that together make up the ciliary architecture [43]. It may be that some aspects of multicilogenesis in this location involve unique genes. It will be interesting to apply comparative analysis with data from other murine multiciliated tissues to see if such unique genes exist.

Multiple mouse models are available for the study of OM. These include mice deficient in genes such as *Tgif1, Mecom*/*Evi1, Fbxo11, Tlr4, Eda* and *Edar [30-,32, 44, 45].* These genes are all expressed in the ALI cultures, although not all of them show expression associated with differentiation. *Mecom/Evi1* is the gene that appears to show the largest difference between the differentiated and undifferentiated cultures. We have recently shown that the *Junbo*^*-/*+^ OM mouse, which is heterozygous for a mutation in *Mecom/Evi1 [32].* develops an exacerbated OM phenotype when *Bpifa1* is also removed [4]. In addition, the ALI cells also exhibit some expression if some known human OM susceptibility genes. Most interestingly, the differentiated cells strongly express BC048546, a presumptive orthologue of A2ML1, a well validated human OM susceptibility gene, that is present in the murine middle ear [29]. Our data suggests that the differentiated mMEEC culture system might have utility for reproducing the OM phenotype of these mouse mutants *in vitro* and enable comparative studies between unaffected and diseased cultures. Our data can also be used to understand the expression and localisation of other disease-causing mutations, for example in genes associated with primary ciliary dyskinesia, an autosomal recessive genetic disorder that causes defects in the action of cilia and results in OM [46].

In conclusion we have presented data describing the gene expression changes in primary murine middle ear cells as they become differentiated when cultured at an ALI. Consistent with our established understanding of the cellular composition of the middle ear this signature describes a mucociliary epithelium. This data set provides a complete transcriptome of the middle ear epithelium that will be a valuable tool for understanding the role played by candidate genes in the middle ear and during the development of OM.

## Author contribution

C.D.B and L.B. designed the study.

A.M., M.M.K.C. and CJ performed the experimental work.

C.D.B. and M.M.K.C. analysed the data.

C.D.B. wrote the paper.

All authors reviewed the paper.

## Acknowledgements

We acknowledge the significant contributions made to establishing the cell model by Professor Steve Brown and Professor Michael Cheeseman at MRC Harwell and thank Dr Paul Heath and Wenbin Wei, at Sheffield Institute for Translational Neuroscience (SITraN), University of Sheffield, for running the arrays and performing the bioinformatic analysis.

## Requests for data

The authors will make the supplementary data and raw array data available on request and it will be deposited prior to formal publication.

